# IDRs lead the way: Cooperativity between intrinsically disordered regions and structured interaction domains drives enrichment in transcription condensates

**DOI:** 10.64898/2026.05.25.727758

**Authors:** Arad Abazari, Dinitha Y. Caldera, Alisha Budhathoki, Ganesh Pandey, Mehdi Shafiei Aporvari, Filmon A. Medhanie, Jan-Hendrik Spille

## Abstract

Many biomolecular condensates are thought to form through phase separation driven by weak and multivalent, non-stoichiometric interactions between intrinsically disordered protein regions (IDRs). IDRs are abundant in the transcription-related proteome. In vitro, different transcription-related IDRs coalesce into the same droplets, providing support for this IDR-centric paradigm of protein enrichment in transcription condensates in the cell nucleus. But our experiments show that IDRs are not sufficient to account for the degree of enrichment observed for full-length proteins in endogenous transcription condensates. Instead, we find a pattern in which IDRs facilitate engagement of structured interaction domains with a binding substrate. Instead, we find a pattern in which IDRs facilitate engagement of structured interaction domains with a binding substrate. Our results indicate that the role of IDRs in transcription condensates requires further investigation with tools that assess their mode of action in situ. Understanding the role of different protein domains and their interplay will also be important for interpreting biotechnological assays that utilize parts of condensate forming proteins.

## Introduction

The precise spatiotemporal regulation of eukaryotic gene expression relies on the action of transcription factors, coactivators, and RNA polymerase II (Pol II) (Cramer, 2019). Formation of phase-separated biomolecular condensates has been proposed to explain how the dozens of factors involved in transcription regulation cooperate effectively (Hnisz *et al*., 2017). Transcription condensates are dynamic, membraneless compartments thought to form at highly expressed, super enhancer-controlled genes (Cho *et al*., 2018; Sabari, 2020). Major protein components of transcription condensates include Pol II, the Mediator complex, and the coactivator BRD4 (Cho *et al*., 2018; Sabari *et al*., 2018). Transcription factors (Boija *et al*., 2018) and RNA (Henninger *et al*., 2021) may also contribute to condensate formation. All of these protein components have prominent intrinsically disordered regions (IDRs).

The prevailing biophysical model for many biomolecular condensates posits that weak, multivalent interactions between these IDRs drive condensate association (Banani *et al*., 2017). Hence, proteins would only loosely associate with these assemblies and readily move in and out of the condensate phase. Cellular evidence for this form of interaction stems from fluorescence recovery after photobleaching (FRAP) experiments that probe the exchange of molecules with the condensate volume (Cho *et al*., 2018; Sabari *et al*., 2018) but interpretation of FRAP data remains qualitative due to technical constraints (McSwiggen *et al*., 2019). In the context of transcription regulation, MED1, a subunit of the Mediator complex, BRD4 (Sabari *et al*., 2018), as well as the C-terminal domain (CTD) of Pol II (Kwon *et al*., 2013) all harbor long IDRs (**Fig. 1A**) that independently undergo phase separation *in vitro*. Moreover, in binary mixtures the Pol II CTD (Guo *et al*., 2019) and the BRD4 IDR (Sabari *et al*., 2018) partition into the same droplets as the MED1 IDR. However, droplet formation assays with purified proteins *in vitro* are typically driven by high protein concentrations and molecular crowding agents while lacking both the full complexity and physical constraints of the cellular environment. These assays are useful to map out phase diagrams under varying environmental conditions and enable precise characterization of the IDR grammar that modulates interaction strength (Wang *et al*., 2018; Choi, Holehouse and Pappu, 2020; Bremer *et al*., 2022) but it is often unclear whether effects observed in this simplified systems are of importance in the cell.

**Fig. 1.**
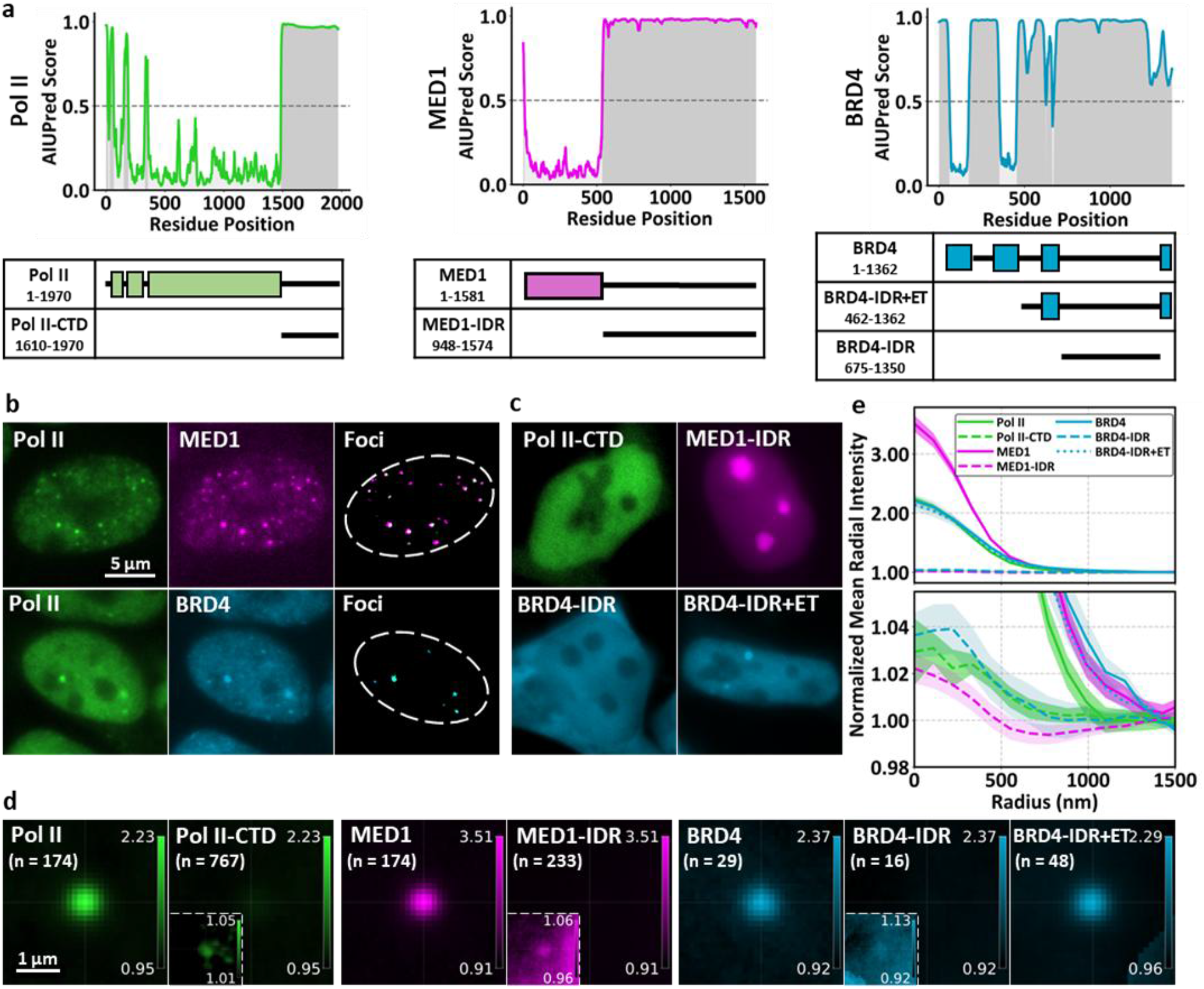
Intrinsically disordered regions (IDRs) are not sufficient to drive enrichment in condensates. **a)** AIUPred disorder probability scores for Pol II (RPB1), MED1, and BRD4. Bottom: Protein domain schematics of the full-length proteins and the truncated constructs used in this study. **b)** Representative widefield fluorescence images of endogenous Snap-Pol II, Halo-MED1, and Halo-BRD4 show colocalization in transcription condensate foci. **c)** In contrast to full-length proteins, the Pol II CTD, MED1 IDR, and BRD4 IDR constructs yield homogeneously distributed signal throughout the nucleus. The MED1 IDR accumulates in, whereas Pol II CTD and BRD IDR are excluded from nucleoli. The BRD4 IDR is also present in the cytoplasm. A BRD4 IDR+ET construct is nuclear localized and forms distinct foci that colocalize with Snap-Pol II condensates. **d)** Aggregate signal at Snap-Pol II condensate centroids for endogenous full-length proteins and exogenous truncation mutants. Number of condensates evaluated: Pol II, n = 174; Pol II CTD, n = 767; MED1, n = 174; MED1 IDR, n = 233; BRD4, n = 29; BRD4 IDR, n = 16; BRD4 IDR+ET, n = 48. **e**) Radial intensity distribution around Snap-Pol II condensate centroids normalized to nuclear average (= 1) from data presented in **e**. Bottom: Magnified view to illustrate low levels of IDR enrichment at endogenous condensates.

We set out to determine how the IDRs used in droplet formation assays – Pol II CTD, MED1 IDR, and BRD4 IDR – would interact with endogenous transcription condensates *in vivo* and whether they are sufficient for the observed enrichment in condensate foci. To address this question, we expressed exogenous fluorescently labeled IDR-only constructs in mouse embryonic stem cell lines. The cells were engineered to also express endogenously tagged condensate proteins (Snap-Pol II, Halo-MED1, and Halo-BRD4). Using two-color fluorescence microscopy, we quantified the partitioning of endogenous full-length proteins and fluorescently tagged versions of their IDRs into the same endogenous condensates.

## Methods

We used CRISPR/Cas9 gene editing technology to label endogenous RPB1 (Pol II), MED1 (Mediator), and BRD4 proteins in mouse embryonic stem cells with enzyme tags (Halo-tag, Snap-tag) as previously described (Cho *et al*., 2018; Sabari *et al*., 2018; Budhathoki *et al*., 2025). We created single- (Snap-Pol II +/+) and double-edited cell lines (Snap-Pol II +/+ - Halo-Med1 +/+; Snap-Pol II +/+ – Halo-BRD4 +/+). Insertion of fluorescent labels did not impact cell growth or protein function. Homozygous cell lines were grown in 2i media.

We cloned plasmids expressing truncated versions of the three proteins labeled with the same tags or fluorescent proteins: Halo-Pol II-CTD, eGFP-MED1-IDR, and Halo-BRD4-IDR. These recombinant plasmids were transfected into the Snap-Pol II +/+ cell line using Lipofectamine 3000. At 24–48 h post-transfection, live cells expressing endogenous full-length and exogenous IDR constructs were incubated with 200 nM Halo-TMR and 50 nM Snap-JFX650 (Grimm *et al*., 2021) for 20 mins. Cells were then washed and incubated in 2i media for 40 minutes before imaging in L-15 + 15% FBS imaging media.

### RNA-Pol II, Med1 and BRD4 are enriched in endogenous transcription condensates

We first quantified enrichment of full-length proteins in transcription condensates by two-color fluorescence microscopy (**Fig. 1B**). Halo-MED1 and Halo-BRD4, respectively, colocalized with Snap-Pol II in readily identifiable condensate foci as previously described by us and many others (Cho *et al*., 2018; Sabari *et al*., 2018). We used the Laplacian of Gaussian function (Scipy) and manual thresholding to segment condensate regions in Snap-Pol II images. Regions of interest around condensate centroids were used to align signal from both imaging channels (**Fig. 1E**). To account for slight differences in expression level, we normalized intensities at each condensate to the mean intensity in the respective nucleus. Thus, each individual region of interest reports relative enrichment before averaging. We excluded nuclear regions inaccessible to the proteins, i.e. nucleoli and cytoplasmic regions. This approach allows us to quantify partitioning of proteins into endogenous condensates. For quantification we plotted the radial intensity profile obtained from the average signal at a large number of condensates (**Fig. 1D**). Relative intensities at the origin indicate the fold-enrichment of the respective protein in condensates. Through this approach, we find Pol II and BRD4 to be enriched approximately 2-fold over background in condensates, whereas MED1 is enriched as much as 3-fold.

### IDRs of the same proteins are not sufficient to drive enrichment in condensates

To determine whether the IDRs of the three major condensate constituents Pol II, MED1, and BRD4 are indeed sufficient for recruiting these proteins to condensates we transfected fluorescently labeled, truncated protein domains into Snap-Pol II cells (**Fig. 1A**). The amino acid sequences of the IDR constructs used here mirror those used in published works for in vitro droplet formation assays (Kwon *et al*., 2013; Sabari *et al*., 2018; Guo *et al*., 2019). We confirmed the coordinates with the AIUPred v2 disorder probability predictor (Erdős and Dosztányi, 2024) (**Fig. 1A**).

After transfection, cells were labeled as described above (**Fig. 1C**). Both the Pol II CTD and the MED1 IDR were predominantly localized to the cell nucleus. The BRD4 IDR was slightly enriched in the nucleus but also distributed throughout the cytoplasm. The MED1 IDR prominently enriches in nucleoli as observed in other studies (Lyons *et al*., 2023; Lee *et al*., 2024). The Pol II CTD and BRD4-IDR were excluded from nucleoli.

To our surprise, neither of the three exogenous constructs formed the prominent foci that we observed for their full-length counterparts. We identified condensate coordinates from Snap-Pol II images acquired in the same nuclei and repeated the enrichment analysis despite no visual indication of IDR accumulation in these endogenous condensates. In the aggregate analysis, we were able to detect a small degree of enrichment over background (**Fig. 1E**). Quantification by radial intensity distribution (**Fig. 1D**) reveals only 1.02 – 1.04-fold enrichment of the IDRs, whereas full-length proteins enriched 2 – 3-fold by the same metric. We confirmed that this was not a result of expression level differences. We note that data acquired by others shows similar findings for the Pol II CTD and MED1 IDR (Lee *et al*., 2024). This unexpected outcome suggests that contrary to the IDR-centric transcription condensate model, the IDRs of these core constituents are not sufficient to explain the enrichment of full-length proteins in condensates.

### Fusing binding domains to IDRs rescues condensate enrichment

We showed previously that in the case of Pol II, the CTD is not sufficient but required for Pol II enrichment in condensates (Budhathoki *et al*., 2025). Neither the CTD by itself, nor a CTD-deletion mutant enriches strongly in condensates. A recent structural biology study proposes that the CTD facilitates Pol II loading onto DNA by anchoring the pre-initiation complex (Zhang *et al*., 2026). Rather than accumulating Pol II through liquid-liquid phase separation, the CTD guides Pol II to access stoichiometric binding sites on DNA. This model is fully consistent with our observations. We also reported that Pol II in condensates is in a Serine 5-phosphorylated state, which reflects initiating and early transcribing Pol II and therefore a chromatin-bound state (Budhathoki *et al*., 2025). In separate studies we show that condensates form at promoter chromatin hubs (Pandey *et al*., 2026; Bogdanović *et al*., 2026). All of these findings emphasize that binding to a chromatin substrate is required for Pol II to enrich in transcription condensates to the degree that we observed for the full-length protein.

A similar effect is at play in the case of BRD4. BRD4 consists of two N-terminal acetyl-chromatin binding bromodomains. The bromodomains allow BRD4 to bind to acetylated histone tails found in active chromatin. The bromodomains are followed by a patch that contains a phosphorylation site and basic interaction domains (BID, aa 484-579), an extra-terminal domain (ET, aa 600-678), a prominent C-terminal IDR (aa 679-1350), and a short C-terminal motif (aa 1351-1362) (Devaiah, Gegonne and Singer, 2016). The ET domain and CTM are known to directly interact with other transcription factors and coactivators (Rahman *et al*., 2011; Wu *et al*., 2013; Zheng *et al*., 2023).

It is important to clarify that early droplet formation assays used specifically the BRD4 IDR (aa 675-1350). This stretch was sufficient for droplet formation by itself and coalesced into MED1 IDR droplets *in vitro* (Sabari *et al*., 2018). We used the same sequence (BRD4 IDR, aa 675-1350) to show that just like the Pol II CTD, the BRD4 IDR is not sufficient for condensate enrichment *in vivo* (**Fig. 1C-E**).

A number of biotechnological applications make use of the condensate forming capability of BRD4 to induce synthetic condensates *in vivo* (Strom *et al*., 2024), coalescence of chromatin loci via condensates (Shin *et al*., 2018) and gene activation (Kim *et al*., 2023), or to determine the proteome of transcription condensates (Lee *et al*., 2024). To focus on effects mediated by BRD4 condensate formation, these applications use a bromodomain deletion mutant frequently termed BRD4-ΔN (aa 442 - 1362), i.e. a combination of BID and ET domain, IDR, and C-terminal motif (aa 441 - 1362). The important distinction between the BRD4 IDR and the N-terminal deletion of the bromodomains (ΔN) is not always made. In some publications BRD4-ΔN is referred to as BRD4-IDR.

To clarify what conveys BRD4 with the ability to enrich in endogenous transcription condensates, we used a similar N-terminal bromodomain deletion mutant (aa 462 – 1362). Like BRD4-ΔN, this BRD4-ET+IDR construct contains the IDR but also the interaction domains flanking it on either side. Repeating the enrichment quantification showed that this modification rescues BRD4 enrichment in condensates. The BRD4-ET+IDR construct forms visible foci that colocalize with and enrich in Snap-Pol II condensates to the same level as full-length BRD4 even after bromodomain deletion (**Fig. 1C-E**).

We conclude that the mechanism of BRD4 enrichment in transcription condensates is in fact fundamentally similar to that of Pol II: A disordered protein domain (Pol II CTD, BRD4 IDR) guides the way to facilitate interactions of a structured interaction domain, which ultimately binds to a substrate to achieve full enrichment. Presence of the interaction domains is required to achieve the degree of enrichment observed for full-length proteins.

We do not know at this point whether the MED1 IDR functions in the same way, but our experiments show that like the Pol II CTD and the BRD4 IDR it is not sufficient to recruit full-length MED1 to endogenous condensates. Among the three IDR constructs, the MED1 IDR showed the lowest enrichment in condensates despite the fact that full-length MED1 was most strongly enriched. FRAP data on both MED1 (Sabari *et al*., 2018) and another Mediator complex subunit, MED19 (Cho *et al*., 2018), had already indicated that Mediator has a fast-exchanging fraction of only 60% and a more stably bound fraction of 40% in transcription condensates. This suggests that substrate-binding plays a significant role in Mediator enrichment in condensates. It is not possible to determine from FRAP data alone whether the fast recovery fraction represents IDR-mediated association or substrate binding times on the timescale of the FRAP recovery (McSwiggen *et al*., 2019).

The MED1 IDR has been used to extract associating proteins in pull-down experiments and to mimic the condensate environment in the cell by tethering it to a highly repetitive Lac-array (Lyons *et al*., 2023). A high local concentration is achieved by tethering a strong LacI binding domain to the MED1 IDR and providing it with the Lac-array binding substrate. In these experiments, too, the local enrichment requires substrate binding. This system is highly successful in mapping out amino acid grammars that convey IDRs with mutual affinity and therefore favor partitioning into the same condensates. But one should keep in mind that readouts are limited to interactions with the protein domain that is anchored at the Lac-array at the high local concentration conveyed by the binding array. The concentration of the anchored domain can far exceed the concentration observed at endogenous condensates, and the array lacks the binding substrate that may ultimately contribute more to enrichment of full-length proteins in endogenous condensates than the IDR interactions themselves.

### A revised model for protein enrichment in transcription condensates

Based on the experiments presented in **Fig. 1** and evidence from literature including our own most recent results (Budhathoki *et al*., 2025; Pandey *et al*., 2026), we propose an updated paradigm for how transcription-related proteins co-enriched in condensate foci in the cell. Prevalent descriptions emphasize phase-separation of disordered protein domains as the mechanism underlying the visible phenotype. Accordingly, schematic representations of transcription condensates typically involve a cloud – or *in vivo* droplet – of loosely associated proteins around a chromatin locus (**Fig. 2A**). This model description leans heavily on results from *in vitro* droplet formation assays. These assays enable the construction of full phase diagrams since sequence composition and environmental parameters (salt, PH, temperature, concentration) can be varied easily in a way that would not be feasible in the cellular context. Droplet formation assays clearly demonstrate that IDRs of condensate-forming proteins have self- and mutual affinities that favor coalesce into the same droplets in binary mixtures (**Fig. 2B**). These interactions carry over in principle to the cellular context, but their strength is modulated by environmental conditions in that native environment. Our experiments show that the focus on IDR interactions is insufficient to explain the presence of transcription condensates. The IDRs of transcription condensate proteins do enrich in endogenous condensates, but to a very minor degree. Structured binding domains are required in all cases to accumulate the transcription machinery (**Fig. 2C**).

**Fig. 2.**
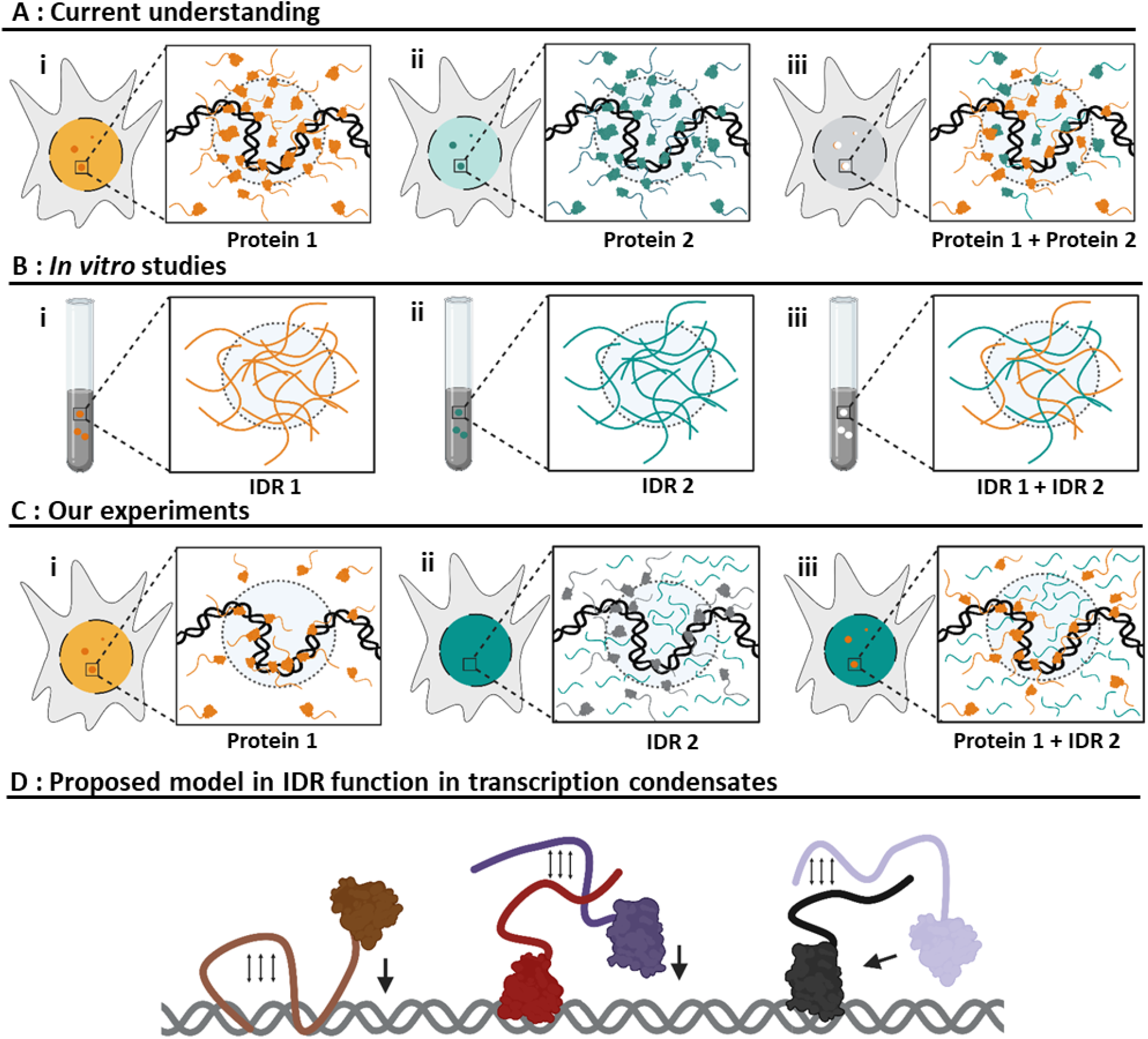
A revised model for the role of IDRs in protein recruitment to transcription condensates. **a)** Fluorescence microscopy of Pol II, MED1, and BRD4 (i, ii) reveals foci in the cell nucleus. (iii) Endogenous proteins coenrich in transcription condensates. Enrichment is thought to result from IDR interactions. **b)** This prevailing model is motivated by the fact that *in vitro*, the Pol II CTD, as well as the MED1 IDR and BRD 4 IDR each form droplets by phase separation (i & ii). Moreover, in binary mixtures these domains coalesce into the same droplets, mirroring the cellular phenotype of full-length proteins. **c)** Our experiments presented in this manuscript reveal that these three domains are insufficient for condensate recruitment of their full-length counterparts to the degree observed in the cell. (i) Full-length proteins form condensate foci, whereas IDR constructs distribute homogenously throughout the nucleus (ii). (iii) Aggregate analysis of dozens of full-length protein condensates reveals only minimal levels of IDR enrichment. **d)** The role of IDRs in recruiting Pol II, MED1, and BRD4 to endogenous transcription condensates is likely more akin to that of disordered transcription factor activation domains that through weak interactions with chromatin or chromatin-bound proteins guide the protein to favored binding substrates (DNA or DNA-bound proteins). Schematic created in BioRender. Caldera, D. (2026) https://BioRender.com/7h49zfs

That is not to say that IDRs do not play a role in the process. For the Pol II we and others have shown that the CTD facilitates substrate binding of the structured domain. We believe that a more accurate model description of the interactions that allow transcription machinery to accumulate in dense condensate foci needs to consider the fact that the weak, transient IDR interactions with chromatin itself, with RNA, or with other proteins likely serve to guide the way for their structured counterparts to more efficiently bind to a local substrate (**Fig. 2D**).

## Conclusions

Intrinsically disordered protein domains have emerged as powerful regulators of biological function (Holehouse and Kragelund, 2024). But IDRs typically do not exist in isolation. We have shown here that the IDRs themselves are not sufficient to explain protein enrichment in transcription condensates to the degree observed for full-length proteins despite their ability to coalesce in droplets *in vitro*. Similar mechanisms as those observed here have been described more thoroughly for transcription factors (TFs). It is well-documented that disordered activation domains of certain TFs modulate affinity for DNA binding sites by favoring interaction with those sites (Már, Nitsenko and Heidarsson, 2023; Jonas, Navon and Barkai, 2025). The IDRs can slide along DNA or interact with chromatin-bound proteins to hold the binding domain in place just long enough to favor targeting to a nearby binding substrate (Morin *et al*., 2022). In contrast, condensate formation by transcription factors with strong activation domains did not increase transcriptional output (Trojanowski *et al*., 2022). All evidence suggests that the three transcription condensate proteins described here, Pol II, MED1, and BRD4, act in a similar manner. The result is a classical biophysical mechanism: Cooperativity between the IDR and the binding domain that enhances binding through additional favorable interactions with adjacent sites. The role of the IDR in this model is to favor selection of or access to specific substrate binding sites, not so much to hold the protein in place within the condensate.

Proteins that harbor structured binding domains alongside IDRs exist on a continuous spectrum of behaviors. The chromatin remodeler cBAF illustrates the other end of the scale. Its ARID1A/B IDRs modulate interactions with other proteins and control positioning on chromatin, but accumulate in very prominent, large condensates upon disruption of the DNA binding domain (Patil *et al*., 2023). In this case, the IDRs have sufficient affinity to drive condensate formation off chromatin. Without perturbation, the binding of the structured domain to the abundant DNA substrate competes against the condensate-forming tendency and titrates away free cBAF. The coactivator BRD4 presents an even more complex case since it harbors several interaction domains that allow it to interact with different substrates: Not only two bromodomains but also additional protein interaction motifs (ET, CTM). The lack of visible puncta of the BRD4 IDR construct indicates that condensate formation of the bromodomain deletion mutant BRD4-ΔN (Strom et al. 2024) likely reflects substrate binding of the ET or CTM domains and not IDR phase separation.

More broadly, a different aspect of the interplay between disordered and structured protein domains is gaining increasing attention. Besides the well-characterized stable binding domain structures discussed here, interaction domains can also form transiently and context-dependent within otherwise disordered regions. These domains can nevertheless result in strong substrate binding (Borcherds *et al*., 2014; Crabtree *et al*., 2017; Wicky, Shammas and Clarke, 2017; Holehouse and Kragelund, 2024; Hess and Joseph, 2025).

The difficulty of studying biomolecular condensates in their cellular context is exacerbated for small foci like transcription condensates (Pandey, Budhathoki and Spille, 2023). Commonly used assays to probe material properties and biophysical mechanisms cannot be interpreted quantitatively if the condensates are diffraction-limited (McSwiggen *et al*., 2019). Replacing endogenous full-length proteins with mutated versions is also often not possible in the cell or would require laborious acute knockdown of essential proteins. Our experiments emphasize the need to replicate reconstitution experiments as closely as possible in the native context to further our understanding of biomolecular condensates structure and function.

## Acknowledgements

We thank Luke D. Lavis, Jonathan B. Grimm, and the Open Chemistry team (Janelia Research Campus) for the gift of Janelia Fluor dyes. We acknowledge support from the Research Corporation for Scientific Advancement (CMC 28407), NIH (R35GM150560), NSF (2306187) and the University of Illinois Chicago Startup Fund. This research was supported in part by grants from the NSF (DMS-2235451) and Simons Foundation (MP-TMPS-00005320) to the NSF-Simons National Institute for Theory and Mathematics in Biology (NITMB).

